# Regulation of islet function and gene expression by prolactin during pregnancy

**DOI:** 10.1101/830836

**Authors:** Vipul Shrivastava, Megan Lee, Marle Pretorius, Guneet Makkar, Carol Huang

## Abstract

Pancreatic islets adapt to insulin resistance of pregnancy by up regulating β-cell proliferation and increase insulin secretion. Previously, we found that prolactin receptor (Prlr) signaling is important for this process, as heterozygous prolactin receptor-null (Prlr^+/−^) mice are glucose intolerant, had a lower number of β cells and lower serum insulin levels than wild type mice during pregnancy. However, since Prlr expression is ubiquitous, to determine its β-cell specific effects, we generated a transgenic mouse with a floxed Prlr allele under the control of an inducible promoter, allowing conditional deletion of Prlr from β cells in adult mice. In this study, we found that β-cell-specific Prlr reduction resulted in elevated blood glucose during pregnancy. Similar to our previous finding in mouse with global Prlr reduction, β-cell-specific Prlr loss led to a lower β-cell mass and a lower in vivo insulin level during pregnancy. However, these islets do not have an intrinsic insulin secretion defect when tested in vitro. Interestingly, when we compared the islet gene expression profile, using islets isolated from mice with global versus β-cell-specific Prlr reduction, we found some important differences in genes that regulate apoptosis and insulin secretion. This suggests that Prlr has both cell-autonomous and non-cell-autonomous effect on β cells, beyond its regulation of pro-proliferative genes.

## Introduction

Pregnancy is characterized by insulin resistance to shunt nutrients from the mother to the developing fetus. Pancreatic β cells adapt to the insulin resistance of pregnancy by up regulating islet mass and insulin secretion, a process that quickly reverses at parturition (25, 31). In animal studies, pregnancy-induced β-cell adaptation consists of 1) a lower threshold for glucose-stimulated insulin secretion, 2) a higher insulin content, and 3) a higher β-cell proliferation rate (5, 22, 23, 30, 33). Both in vitro and in vivo observations support a role for prolactin (PRL) and/or placental lactogens (PLs) in this adaptive process. First, prolactin receptor (Prlr), the receptor for both PRL and PLs, is present on pancreatic β cells and Prlr expression increases during pregnancy (1, 9, 17, 19, 32). Second, the rise in PRL and PLs levels parallel the increases in β-cell mass and glucose-stimulated insulin secretion during pregnancy (29). Third, in vitro exposure of isolated islets to PRL/PLs increases insulin secretion, β-cell proliferation and lowers the threshold of glucose-stimulated insulin secretion (6, 7, 32), mimicking the effects of pregnancy on β cells. To determine whether Prlr signaling is required for β-cell adaptation to pregnancy in vivo, transgenic mice with global or β-cell specific Prlr knockout have been examined. Using heterozygous prolactin receptor-null mice (Prlr^+/−^), we confirmed the in vivo role of Prlr on β-cell adaptation to pregnancy (12). We found that in comparison to their wild type littermates, pregnant Prlr^+/−^ dams were glucose intolerant and secreted less insulin in response to an intraperitoneal glucose tolerance test (IPGTT). They also had a lower β-cell proliferation rate and lower β-cell mass than the Prlr^+/+^ mice. We could not use the homozygous Prlr^−/−^ mice because they have a placental implantation defect, thus are infertile. Two separate groups have since generated β-cell specific Prlr-knockout mice, one using the rat insulin promoter controlled Cre (i.e. RIP-Cre) and the other using the Pdx1 promoter controlled Cre (i.e. Pdx1-Cre)(3, 20, 24). Similar to our findings in Prlr^+/−^ mice, β-cell specific Prlr-deletion caused gestational diabetes, accompanied by a reduction in β-cell proliferation and a blunted glucose-stimulated insulin secretion. However, since both the insulin promoter and the Pdx1 promoter are expressed during embryonic development and they causes a defect in β-cell mass at birth, we generated an inducible conditional Prlr knockout mouse to circumvent this developmental defect. Furthermore, since prolactin receptor is ubiquitously expressed, Prlr may have non-cell autonomous effect on β cells. Our objective is to determine the cell autonomous effect of Prlr during pregnancy, and to explore the non-cell autonomous effect of Prlr by comparing islet gene expression using mice with global and β-cell-specific prolactin receptor deletion.

## Methods

### Ethical approval

All experimental procedures were approved by the Animal Use Review Committee at the University of Calgary in accordance with standards of the Canadian Council on Animal Care.

### Mice

Heterozygous prolactin receptor null mice (Prlr^+/−^) on a C57BL/6 background were purchased from Jackson Laboratory and a working stock was generated by crossing Prlr^+/−^ mice with wild type Prlr^+/+^ mice. The pups were genotyped as previously described (4).

### Generation of the conditional prolactin-receptor null mice

A promoter-driven targeting cassette for the generation of a knockout-first allele with potential for conditional knockout was obtained from EUCOMM (The European Conditional Mouse Mutagenesis Program)(27). The vector was linearized at AsiSI-site and electroporated into mouse ES cells. Neomycin resistant cells were isolated and selected by Southern analysis. ES cells that contain the correctly recombined genome and correct karyotype were then injected into 8-cell CD1E wild type mouse embryos to generate chimeras. The male chimeric mice were then crossed to albino C57BL/6 mice (Charles River, B6N-*Tyr*^C-Brd/BrdCrCrl^) to obtain germline transmission of the mutation. The FRT-flanked neo cassette was removed by crossing with FLPeR mice (The Jackson Laboratories; B6.129S4-*Gt(ROSA)26Sor*^*tm1(FLP1)Dym*^/RainJ). This generated the mice heterozygous for floxed exon5 (*Prlr*^*tm1c*(EUCOMM)Hmgu^, herein denoted as *PrlR*^fl/+)^ that were phenotypically wild-type. It was back-crossed with C57BL/6J mice (The Jackson Laboratory) for more than 10 generations. The PrlR^fl/+^ mice were crossed with Pdx1CreER™ mice (10), which have the *Cre-ER™* gene under the control of the *Pdx1* promoter. The male Pdx1CreER™:PrlR^fl/+^ mice were crossed with female PrlR^fl/+^ to generate the homozygous conditional knockout Pdx1CreER™:PrlR^fl/fl^ (herein denoted as βPrlR^fl/fl^), and the heterozygous conditional knockout βPrlR^fl/+^, and the control littermates, i.e. Pdx1CreER™, PrlR^fl/+^ and PrlR^fl/fl^ mice. The βPrlR^fl/fl^ mice were then crossed with the mT/mG reporter mice, which carries *ROSA*^*mT/mG*^, a cell membrane-targeted, two-color fluorescent Cre-reporter allele. Before Cre recombination, the tdTomato (mT) fluorescence expression is widespread in all cells. Upon Cre recombinase activation, the cells express membrane-localized EGFP (mG) fluorescence, replacing the red fluorescence(18). To induce Cre recombinase activity, tamoxifen (dissolved in corn oil) was given by oral gavage at age 8 weeks for 3 days at a dose of 8mg/day (for a 20g mouse). Mice were used 4 weeks after tamoxifen administration.

Mice were maintained on a 12-h light, 12-h dark cycle with liberal access to food and water. Mice were studied at 3-4 months of age. For pregnancy, female mice were set up with wild type male mice and the morning when a vaginal plug was found was designated as day 1 of pregnancy. Mice were used on days 15 of pregnancy. We chose to study day 15 of pregnancy because β-cell proliferation and β-cell mass peaks on days 14-15 of pregnancy (29).

### Intraperitoneal glucose tolerance test (IPGTT) and insulin tolerance test (ITT)

IPGTT (20% D-glucose solution, 2g/kg body weight) and ITT (NovoRapid insulin, 0.5unit/kg body weight) were performed as previously described (12). Additional blood samples (~30μl) were taken at times 0, 5, and 30 minutes of IPGTT for insulin concentration measurements by ELISA (ALPCO, catalog #80-INSMSU-E01). Non-fasted blood glucose was determined using a glucometer (FastTake) by sampling from tail vein at 8am, and at the same time, an additional 30μl of serum was taken from the saphenous vein and stored at −80°C for later measurement of insulin by ELISA.

### Materials

Collagenase P (Ref#11 213 865 001), trypsin 1mg tablets (T7168) and all chemicals are from Sigma. Sources of antibodies are specified below.

### Immunostaining

Pancreas was isolated from pregnant and non-pregnant mice, weighed, embedded in paraffin blocks, longitudinally serial sectioned to 7 μm then stained for insulin to identify β cells as previously described (12). Briefly, after 1 hour of blocking with 1% goat serum/PBS at room temperature, tissues were incubated with primary antibody over night at 4°C (guinea pig anti-insulin at 1:750, DAKO; diluted in 1% goat serum/PBS). This was followed by 1-hour incubation with fluorophore-conjugated secondary antibodies (Cy3-anti-guinea pig, Jackson Laboratories, diluted in 1% goat serum/PBS at 1:300). Bis-benzimide H 33342 trihydrochloride (0.1μg/ml, Sigma) was added to the secondary antibody for nuclear staining. Stained sections were mounted using DakoCytomation fluorescent mounting medium and stored at 4°C.

### Islet Morphometry

Consecutive images of non-overlapping, adjacent areas of the entire pancreas section were acquired using a Zeiss fluorescence microscope, and captured with a CoolSnap digital camera (12). Images were analyzed by ImageJ software to measure the insulin-positive area as well as the area of the entire pancreas section (identified by nuclear staining). β-cell mass was calculated by multiplying the pancreas weight by the β-cell fraction (i.e. the ratio of insulin-positive cell area to total pancreatic tissue area on the entire section. Results represent the average of 6-8 tissue sections per animal from 5-6 animals from each genotype.

### Islet Isolation

The pancreas was first distended using collagenase P (0.66mg/ml in Hank’s Balanced Salt Solution, 2.5ml/pancreas), surgically removed and then incubated at 37°C for 15min under constant agitation. Islets were hand-picked and cultured overnight in RPMI1640 supplemented with 10% fetal bovine serum and penicillin/streptomycin(11).

### Islet RNA isolation and quantitative real-time RT-PCR

Total islet RNA (100–200 islets/mouse) was extracted using the RNeasy Mini Kit (Qiagen). RNA concentration and integrity were assessed using the ND-1000 Spectrophotometer (NanoDrop). cDNA was synthesized using the Quantitect Reverse Transcription Kit (Qiagen). Reactions were carried out in triplicate with QuantiFast SYBR Green Master Mix (Qiagen) at an annealing temperature of 60°C. Data were collected using the DNA Engine Opticon2 Continuous Fluorescence Detection System (BioRad) and software (Bio-Rad). Primer identifying both long and short forms of prolactin receptor was designed using Primer Designing Software (NCBI); sequence: Forward: 5’ ATCTTTCCACCAGTTCCGGG, and Reverse: 5’ TTGGAGAGCCAGTCT CTAGC. The primer sequence for estrogen receptor 1 (ESR 1) is: Forward: TCTGCCAAGGAGACTCGCTACT, and Reverse: 5’ GGTGCATTGGTTTGTAGC TGGAC. The relative amount of RNA was determined by comparison with phosphoglycerate kinase 1 (Pgk1) mRNA as a reference gene as previously described (11).

### RNAseq analysis

Total RNA (100–200 islets/mouse) was extracted using the RNeasy Mini Kit (Qiagen). RNA quality and quantity was measured on the Agilent 2200 TapeStation System. Only samples with a RNA integrity number above 7 were used. Messenger RNA was enriched and separated from rRNA using the NEBNext^®^Poly(A) mRNA Magnetic Isolation Module (New England Biolabs). RNA library was prepared using the NEBNext^®^ Ultra II Directional RNA Library Prep Kit for Illumina (New England Biolabs). Sequencing is performed on Illumina NextSeq 550 System. Gene network analysis was performed using Qiagen’s Ingenuity Pathway Analysis platform to identify differentially expressed genes.

### Statistical analysis

All statistics were performed using GraphPad Prism 4 software. Two-tailed Student’s t tests or ANOVA with Bonferroni post-tests were performed where appropriate. Comparisons were made between the homozygous conditional knockout (βPrlR^fl/fl^), the heterozygous conditional knockout (βPrlR^fl/+^), the wild type littermate (βPrlR^+/+^), or the Prlr^+/−^ mice, as stated in the Figure Legend.

## Results

### Generation of inducible, pancreas-specific prolactin receptor deleted transgenic mice

To understand the role of prolactin receptor signaling on pancreatic β-cell mass and function during pregnancy, we generated mice harboring an inducible conditional deletion of Prlr from the pancreas. We interbred the Pdx1CreER™ mouse, which has a tamoxifen-inducible Cre recombinase under the control of Pdx1, with PrlR^fl/fi^ mouse, which has a floxed deletion of exon 5 of prolactin receptor, to generate the Pdx1CreER™: PrlR^fl/fl^ mouse, or the βPrlR^fl/fl^ mouse. To allow visual determination of the tissue-specific prolactin receptor deletion, we intercrossed the βPrlR^fl/fl^ mouse with the ROSA^mT/mG^ indicator mouse. The ROSA^mT/mG^ mouse constitutively expresses a tdTomato transgene and upon Cre recombination, GFP become expressed in the islets (Supplemental Figure). To induce Cre recombinase expression, mice were given tamoxifen at age 8-10 weeks. We found that in comparison to control littermate without the floxed Prlr allele, i.e. the βPrlr^+/+^ mice, βPrlR^fl/+^ mice had a ~40% reduction in prolactin receptor expression in isolated islets (Fig. 1). In comparison to their respective wild type littermates, we found similar magnitude of reduction in prolactin receptor expression in the islets of βPrlR^fl/+^ and Prlr^+/−^ mice, the latter are transgenic mice with global heterozygous prolactin receptor deletion (12). The reduction in prolactin receptor is specific to the pancreatic islets, as we observed no significant reduction in the prolactin receptor expression in the hypothalamus, fat, and liver (Fig. 1), tissues that play a role in glucose homeostasis.

**Figure 1.**
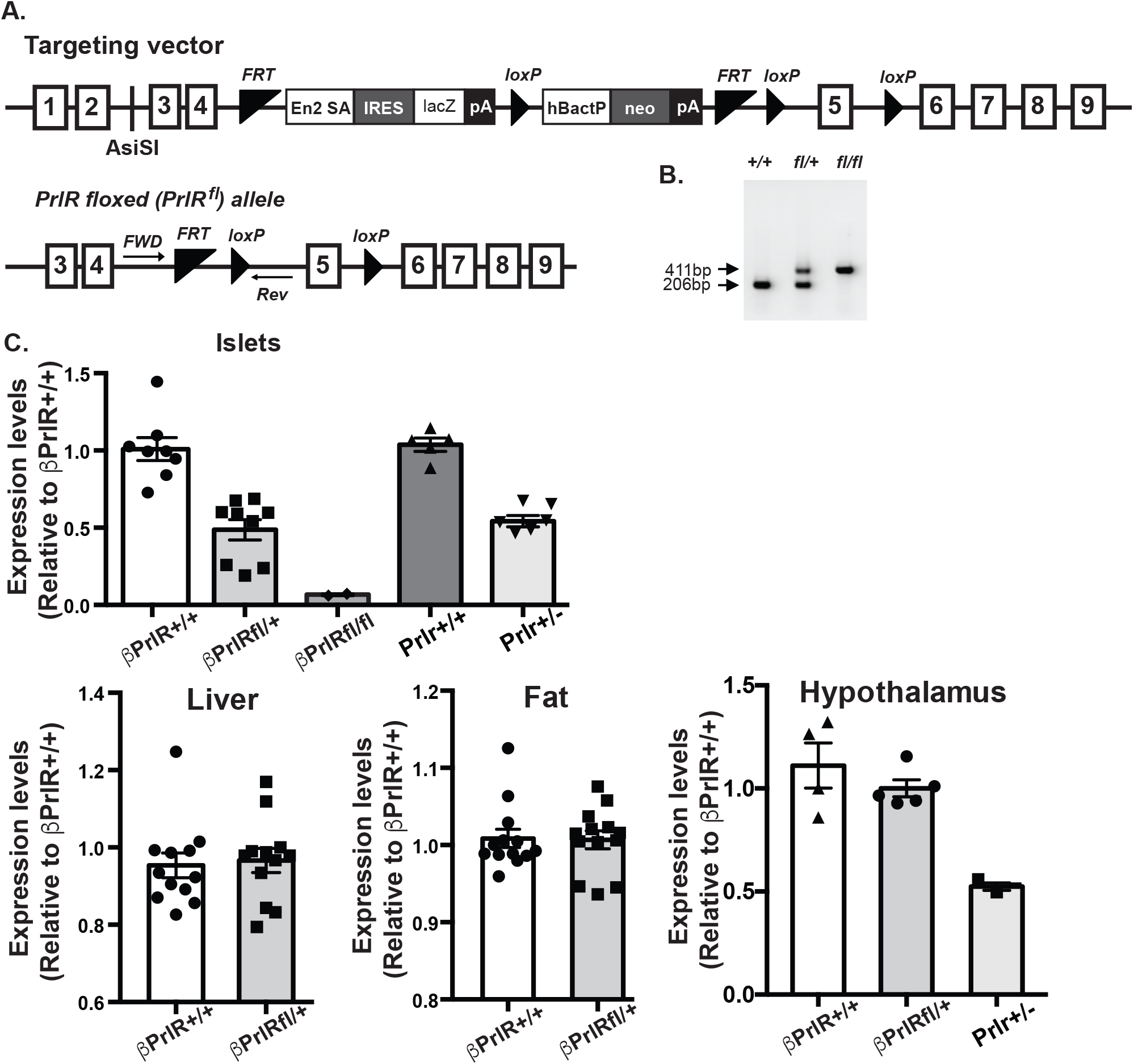
Islet-specfic reduction in Prlr expression of the βPrlrfl/+ mice. Condition deletion of Prlr signaling in pancreatic β cells (βPrlR^fl/fl^). A) Schematic diagram of the targeting vector and floxed PrlR allele. B) Genotyping of a litter that contain mice with homozygous floxed PrlR allele (fl/fl), heterozygous floxed PrlR allele (fl/+), or homozygous wild type (non-floxed) PrlR allele (+/+). C) Relative mRNA expression of prolactin receptor in islets, liver, fat, and hypothalamus. n=3-12 mice/genotype. “*”= p<0.05 and “**”= p<0.01 in comparison to the βPrlR^+/+^ mice.

### Gestational glucose intolerance in βPrlR^fl/+^ mice

Next, we determined glucose homeostasis of the βPrlR^fl/+^ mice during pregnancy. Intraperitoneal glucose tolerance test was performed on day 15 of pregnancy. In comparison to their wild type littermates (βPrlr^+/+^), the βPrlR^fl/+^ mice were glucose intolerant, with greater glucose excursion, expressed as integrated area under the curve over the 120 min of IPGTT (βPrlR^fl/+^:AUC=1869.83±101.21mM × min vs. βPrlr^+/+^:AUC=1324.90± 134.41mM × min, n=10-16)(Fig. 2a-b). There is no significant difference in fasting blood glucose, but random non-fasted blood glucose is higher in the βPrlR^fl/+^ mice (8.14±0.29mM vs. 7.0±0.30mM for βPrlr^+/+^ mice, n=10-16)(Fig. 2c-d). Interestingly, the βPrlR^fl/fl^ mice, which has a 90% reduction in prolactin receptor expression, has a similar degree of glucose intolerance as the βPrlR^fl/+^ mice (Fig. 2). As expected, there is no difference in insulin tolerance between the wild type and mutants (Fig. 2e), as prolactin receptor is only deleted from the pancreatic islets and not from other insulin responsive tissues, such as liver and fat (Fig. 1).

**Figure 2.**
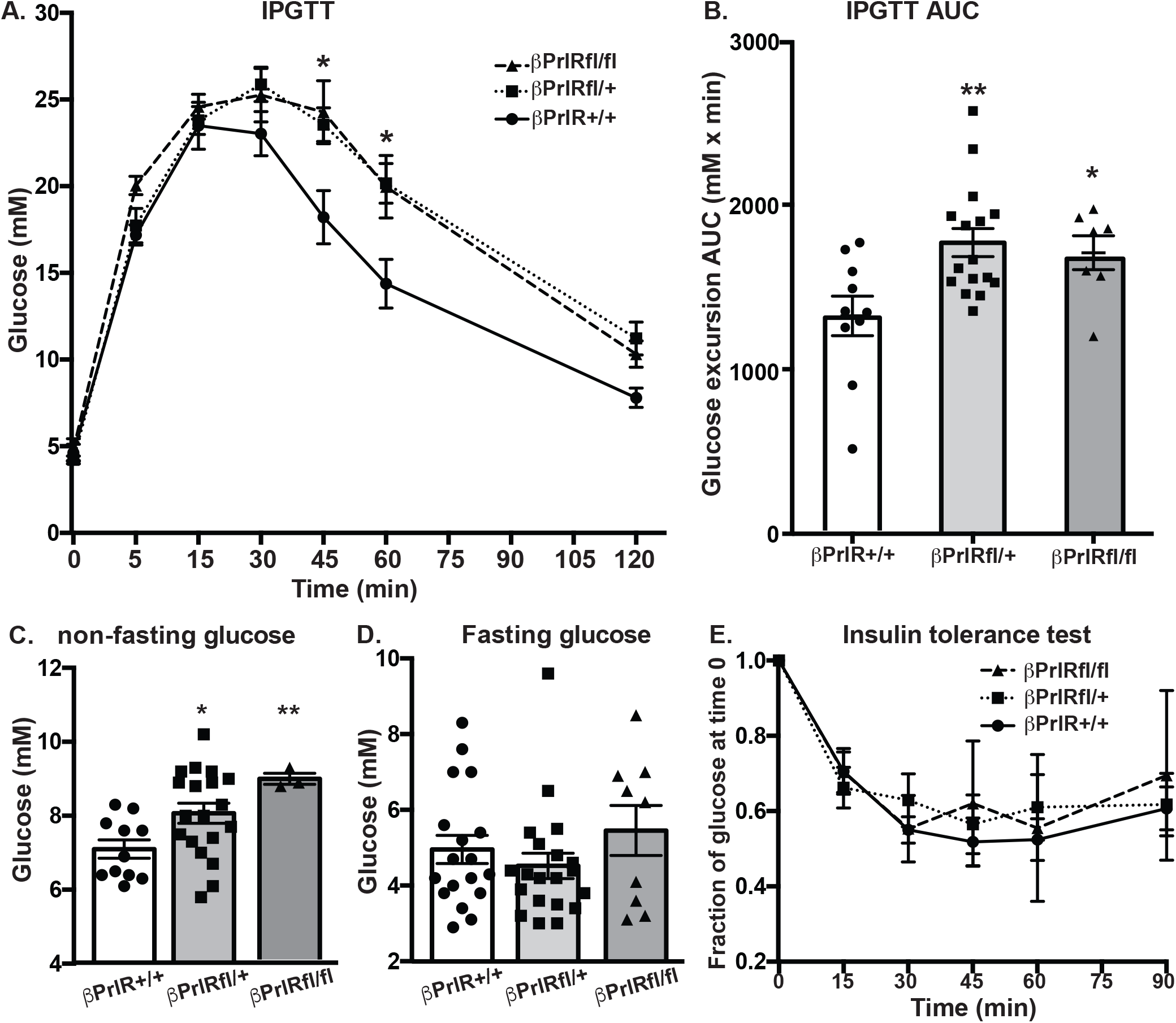
βPrlrfl/+ and βPrlrfl/fl mice are glucose intolerant. βPrlR^fl/+^ and βPrlR^fl/fl^ mice are glucose intolerant during pregnancy. A) Glucose excursion during an intraperitoneal glucose tolerance test. “*”=p<0.05 in comparison to the wild type βPrlR^+/+^ mice at that time point. B) Integrated area under the curve of glucose excursion during an IPGTT. C) Random non-fasting blood glucose. D) Fasting blood glucose. E) Glucose excursion during an intraperitoneal insulin tolerance. Blood glucose are expressed as a fraction of the blood glucose at time For B-E), “*”=p<0.05 and “**”=p<0.01 in comparison to the wild type βPrlR^+/+^ mice. n=3-19 mice per genotype.

### βPrlR^fl/+^ mice had lower β-cell mass and secreted less insulin

Similar to our previous observation in the heterozygous global Prlr-deletion mice (Prlr^+/−^)(12), βPrlR^fl/+^ mice had a lower β-cell mass than their wild type littermates on day 15 of pregnancy (Fig. 3). During an IPGTT, βPrlR^fl/+^ secreted less insulin (expressed as integrated area under the curve)(Fig. 4). Interestingly, the βPrlR^fl/+^ mice had slightly higher fasting insulin, although there is a very wide variability. To determine whether the blunted in vivo insulin secretion is due to a decrease in β-cell mass or the islets are less able to respond to glucose, we performed in vitro glucose-stimulated insulin secretion test. Here, we found that islets from βPrlR^+/+^ and βPrlR^fl/+^ mice were equally responsive to glucose, suggesting that the difference in in vivo insulin secretion was a result of the smaller β-cell mass of the βPrlR^fl/+^ mice.

**Figure 3.**
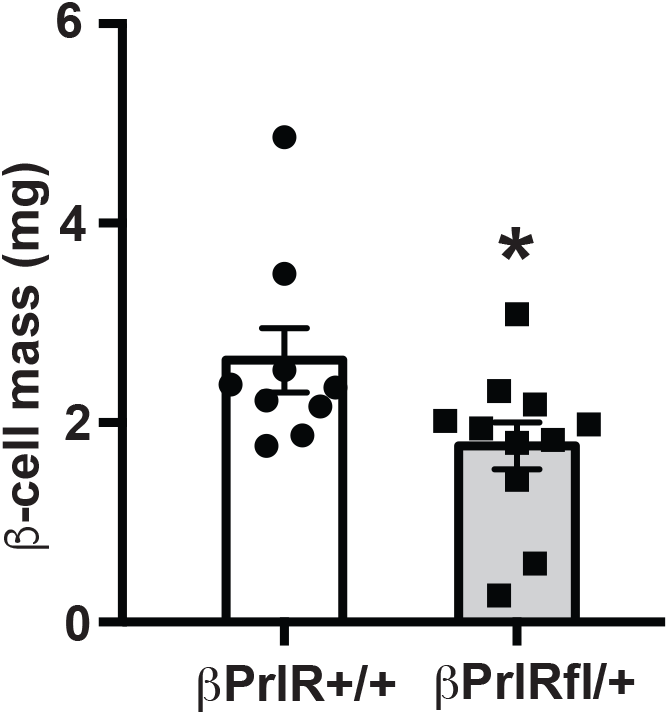
β-cell mass. βPrlR^fl/+^ mice have a lower β-cell mass on day 15 of pregnancy in comparison to the wild type βPrlR^+/+^ mice. “*”=p<0.05. n=9-10 mice per genotype.

**Figure 4.**
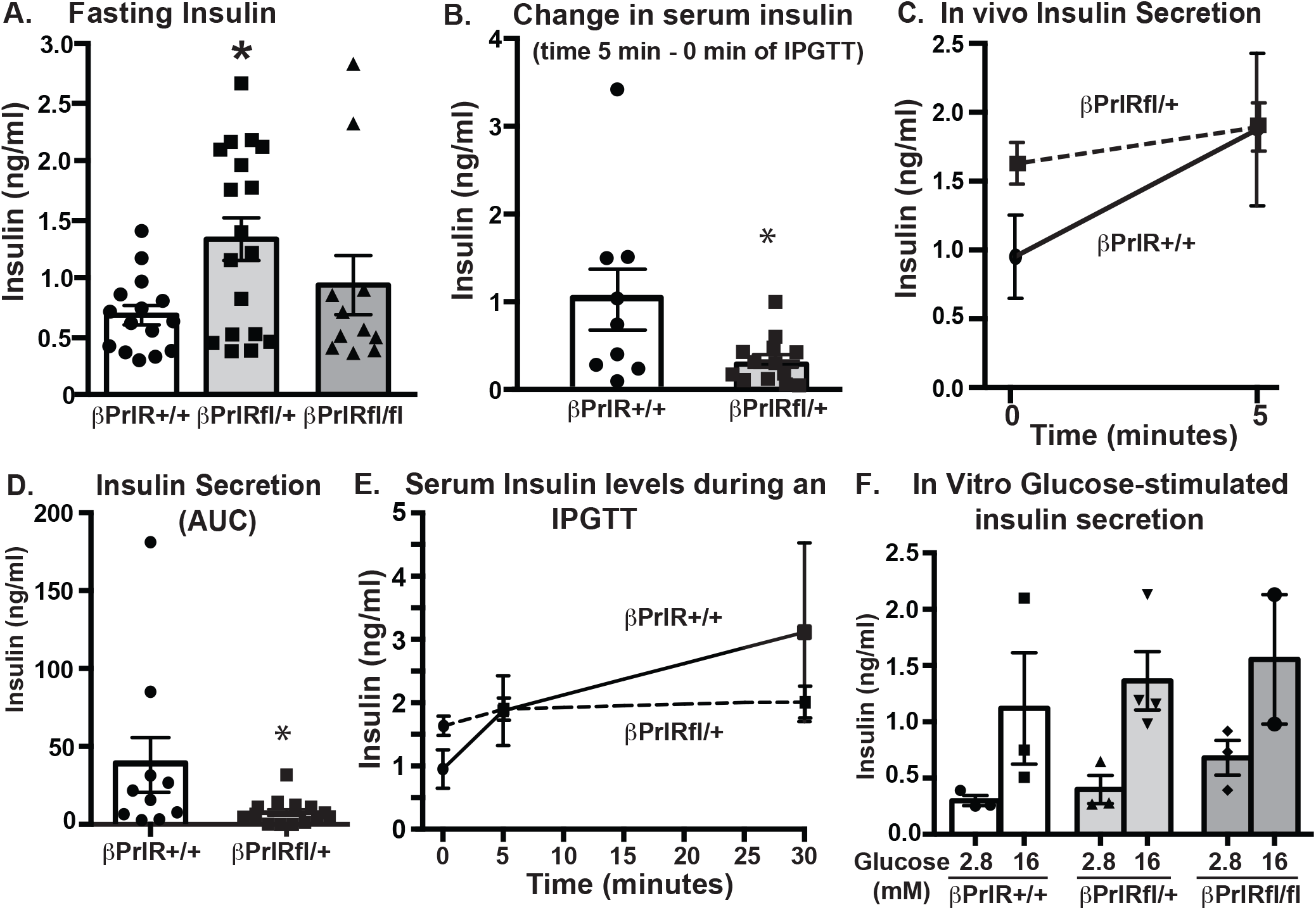
βPrlr^fl/+^ mice secreted less insulin during an IPGTT. βPrlR^fl/−^ mice secreted less insulin during an IPGTT in comparison to the wild type βPrlR^+/+^ mice. A) Fasting insulin. B) Increase in insulin levels during the first 5 minutes of the IPGTT. “*”=p<0.05 in comparison to the βPrlR^+/+^ mice. C) Insulin levels during the first 5 minutes of the IPGTT. D) Integrated insulin secretion during the first 30 minutes of the IPGTT, expressed as area under the curve. “*”=p<0.05 in comparison to the βPrlR^+/+^ mice. E) Serum insulin levels during the first 30 minutes of the IPGTT. F) In vitro glucose-stimulated insulin secretion. n=10-12 mice/genotype for in vivo insulin secretion. n=3 mice/genotype for in vitro insulin secretion.

### Prolactin receptor has non-cell autonomous effects on β-cell gene expression during pregnancy

The above results demonstrated that transgenic mice with global (Prlr^+/−^) or β-cell specific prolactin receptor deletion (βPrlR^fl/+^) are phenotypically similar: both are glucose intolerant on day 15 of pregnancy, with lower β-cell mass and secreted less insulin in response to glucose, in comparison to their wild type littermates. However, since prolactin receptor is ubiquitously expressed, including on the endothelial and epithelial cells in islets, we want to compare the gene expression profile of islets from βPrlR^fl/+^ to that of the Prlr^+/−^ mice, to determine whether a reduction in Prlr in non-β cells can secondarily affect β-cell gene expression. We performed RNAseq analysis and found that estrogen receptor 1 expression was lower in the Prlr^+/−^ islets in comparison to βPrlR^fl/+^ islets (Fig 5). Ingenuity pathway analysis found several canonical pathways that are differentially expressed, including the serotonin receptor-signaling pathway, which has been previously found to regulate β-cell proliferation during pregnancy (13).

**Figure 5.**
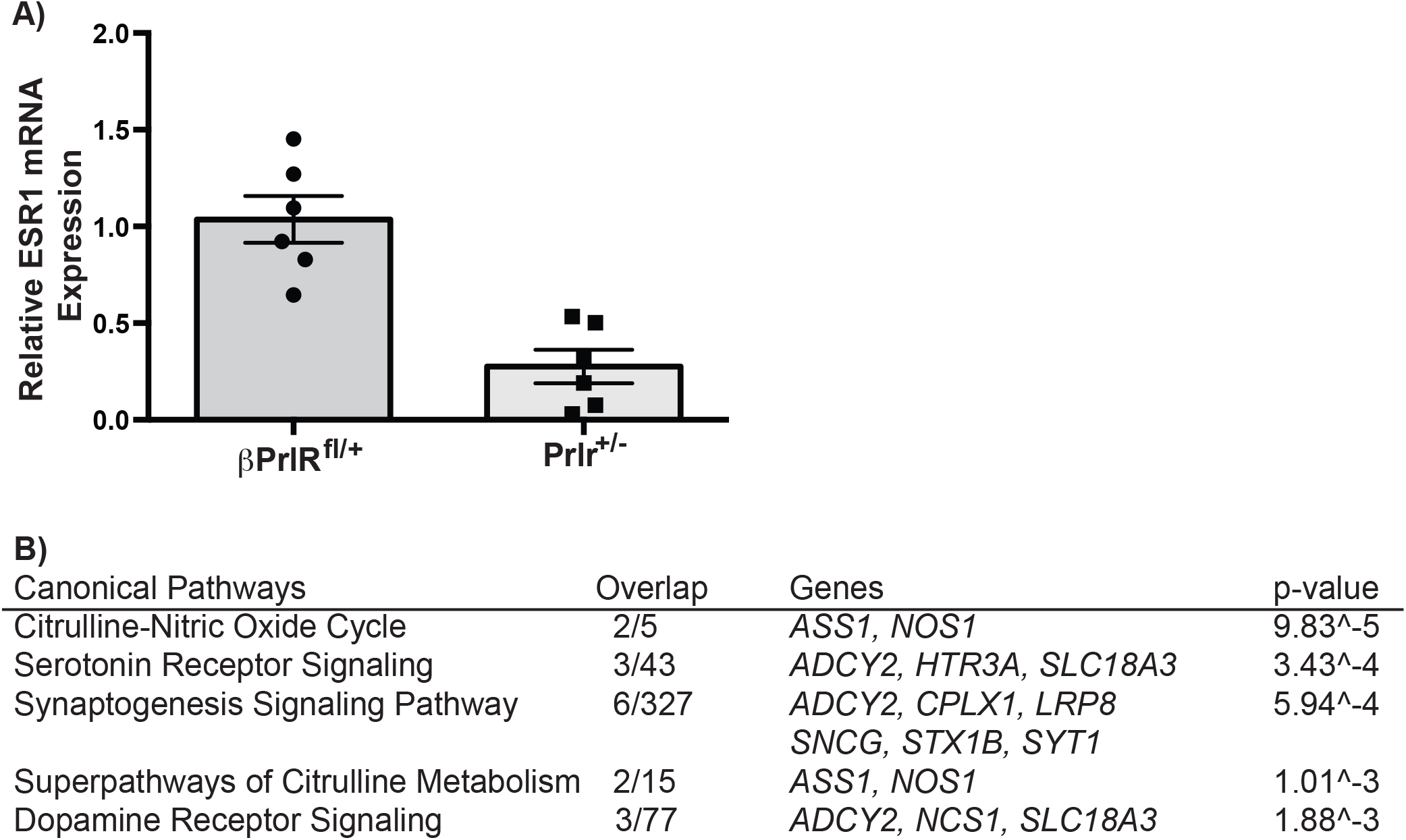
Comparison of gene expression between βPrlRfl/+ and Prlr+/− mice. Differential gene expression between the global (Prlr^+/−^) and β-cell-specific (βPrlR^+/−^) prolactin receptor deleted mice. A) Estrogen receptor 1 expression is significantly lower in the Prlr^+/−^ mice. b) Most significant canonical pathways across the entire dataset.

## Discussion

Pregnancy is an insulin resistant state and maternal pancreatic islets adapt to the increase in insulin demand by up regulating β-cell proliferation and increase insulin secretion. Previously, we have shown that a transgenic mouse with global, heterozygous deletion of prolactin receptor (Prlr^+/−^) was glucose intolerant during pregnancy. This was accompanied by a reduction in β-cell proliferation, β-cell mass, and a blunted glucose-stimulated insulin secretion in comparison to their wild type littermates (12). A caveat of using a global knockout mouse is that the ubiquitous expression of prolactin receptor raises the possibility that the phenotype observed in the pancreatic islet is secondary to prolactin action in tissues other than β cells. To address this possibility, we generated a transgenic mouse with an inducible, conditional prolactin receptor gene knockout. We chose to make the gene deletion inducible because β-cell specific promoters such as the insulin promoter or the Pdx1 promoter are expressed during embryonic development of the pancreas (26). In fact, Auffret et al found that transgenic mouse with a global, homozygous deletion of prolactin receptor (i.e. Prlr^−/−^) has abnormal β-cell mass expansion both embryonically and during perinatal life (2). They found a 30% reduction in β-cell mass in the Prlr^−/−^ newborns. Hence, an inducible gene deletion during adulthood would circumvent this developmental defect and allow us to study prolactin receptor effect specifically during pregnancy.

Several groups have since generated conditional knockout of the prolactin receptor gene. Banerjee et al used a RIP-Cre promoter to generate a conditional Prlr knockout (3). They found that β-cell specific deletion of Prlr led to gestational diabetes due to reduced β-cell proliferation and failure to expand β-cell mas during pregnancy. They identified MafB as one of the Prlr-signaling target, and MafB deletion is maternal β cells caused gestational diabetes as well. Subsequently, they performed a transcriptome analysis and found that forkhead box protein M1, polycomb repressor complex 2 subunits, Suz12 and enhancer of zeste homolog 2 are Prlr signaling targets (24). Interestingly, Prlr was not required for β-cell adaptation to high fat feeding. In fact, pregnancy and high fat feeding activate very different genes in the islets, suggesting that the two metabolic stressors engage different mechanisms to adapt. Interestingly, RIP-Cre is expressed in the arcuate nucleus(28) and Prlr has been specifically deleted from the arcuate nucleus but the effect on glucose homeostasis was not reported(8). Nteeba et al also generated a β-cell-specific Prlr knockout mouse, using a Pdx1-Cre promoter(20). Consistent with Banerjee et al., they found glucose intolerance during second pregnancy. Interestingly, the β-cell-specific Prlr knockout mice had lower β-cell mass in non-pregnant state and the pregnancy-stimulated β-cell mass expansion was essentially non-existent in their model. Moreover, they found that the loss of maternal pancreatic Prlr-signaling was association with a reduction in body weight of the offspring, at embryonic day 15 and on day 1 after delivery. There is also significant dysregulation of placental prolactin family of genes.

Prolactin receptor expression is ubiquitous, and it is expressed on the endothelial cells and ductal cells in the islets. To identify potential non-autonomous effect of Prlr on β cells, we performed RNAseq analysis, comparing gene expression of islets from Prlr^+/−^ vs. βPrlR^fl/+^ mice on day 15 of pregnancy. There is a reduction in estrogen receptor 1 (ESR1) expression in the Prlr^+/−^ mice; this is interesting because estrogen receptor protect β cells from apoptosis (15). Our previous observation that islets from Prlr^+/−^ mice are more sensitive to free fatty acid induced apoptosis than islets from Prlr^+/+^ mice (16) may be in part attributable to this down regulation of ESR1 in islets of Prlr^+/−^ mice. Another interesting finding from our RNAseq analysis is that Ingenuity pathway analysis identified serotonin receptor signaling as one of the top canonical pathway that is differentially regulated between Prlr^+/−^ and βPrlR^fl/+^ mice (13, 21). Kim et al previously reported that serotonin acts downstream of Prlr signaling to stimulate β-cell proliferation, and this was through the Gaq-linked serotonin receptor 5-hydroxytryptamine receptor –2b (Htr2b)(13). In a subsequent study, they found that another serotonin receptor, Htr3a is important for regulation of insulin secretion(14). Taken together, the results from the 3 different β-cell-specific Prlr knockout mice and the global Prlr knockout mice support that Prlr has cell autonomous and non-cell autonomous effect on β-cell adaptation to pregnancy, with the cell autonomous effect accountable for most of the observed effect on β-cell mass expansion and insulin secretion.

## Acknowledgements

We thank Ken Ito and Yaping Yu for technical assistance in the generation of the βPrlR^fl/fl^ mice.

## Author Contributions

CH conceived and designed the project. VS, ML, MP, GM and CH performed experiments and analyzed the data. CH drafted the manuscript. All authors reviewed and approved the manuscript.

---

**Supplemental Figure.**
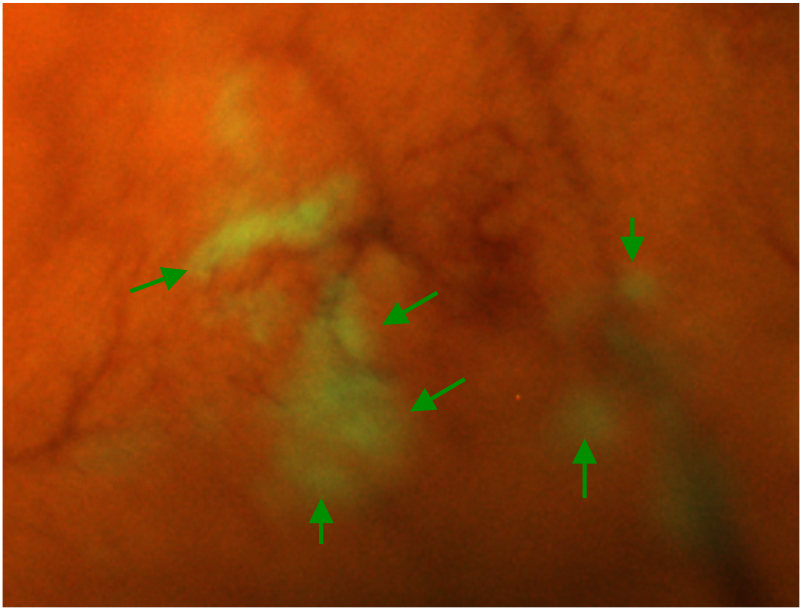
βPrlR^fl/fl^ mice expresses mT/mG, which allow detection of Cre recombination, visualized as green fluorescence (green arrows). The areas of green fluorescence represent islets.

